# What is an urban bird? Trait-based predictions of urban tolerance for the world’s birds are modulated by latitude and human population density

**DOI:** 10.1101/2022.10.28.514262

**Authors:** Montague H. C. Neate-Clegg, Benjamin A. Tonelli, Casey Youngflesh, Joanna X. Wu, Graham A. Montgomery, Çağan H. Şekercioğlu, Morgan W. Tingley

**Affiliations:** Ecology and Evolutionary Biology, University of California, Los Angeles, CA 90095, USA; Ecology, Evolution, and Behavior Program, Michigan State University, East Lansing, MI 48824, USA; School of Biological Sciences, University of Utah, Salt Lake City, UT 84112, USA; Faculty of Sciences, Koç University, Rumelifeneri, Istanbul, Sarıyer, Turkey

**Keywords:** body mass, citizen science, clutch size, diet breadth, dispersal ability, eBird, habitat breadth, longevity, NDVI, nest type

## Abstract

As human density increases, biodiversity must increasingly co-exist with urbanization or face local extinction. Tolerance of urban areas has been linked to numerous functional traits, yet few globally-consistent patterns have emerged to explain variation in urban tolerance, which stymies attempts at a generalizable predictive framework. Here, we calculate an Urban Association Index (UAI) for 3768 bird species in 137 cities across all permanently inhabited continents. We then assess how UAI varies as a function of ten species-specific traits and further test whether the strength of trait relationships vary as a function of three city-specific variables. Of the ten species traits, nine were significantly associated with urban tolerance. Urban-associated species tend to be smaller, less territorial, have greater dispersal ability, broader dietary and habitat niches, larger clutch sizes, greater longevity, and have lower elevational limits. Only bill shape showed no global association with urban tolerance. Additionally, the strength of several trait relationships varied across cities as a function of latitude and/or human population density. For example, the effects of body mass and diet breadth are more pronounced at higher latitudes, while the effects of territoriality and longevity were reduced in cities with higher population density. Thus, the importance of trait filters in birds varies predictably across cities, indicating biogeographic variation in selection for urban tolerance that could explain prior challenges in the search for global patterns. A globally-informed framework that predicts urban tolerance will be integral to conservation as increasing proportions of the world’s biodiversity are impacted by urbanization.

## Introduction

Urbanization is one of the greatest threats to biodiversity (Mcdonald *et al*. 2008; McKinney 2008; McDonald *et al*. 2020). By 2030, an expected 5.2 billion people will live in urban areas alone (United Nations 2018) and urban land cover is predicted to exceed 1.2 million km^2^ globally (Seto *et al*. 2012). Urbanization is accompanied by a consistent loss of biodiversity (Aronson *et al*. 2014; Evans *et al*. 2018; de Camargo Barbosa *et al*. 2020), including reduced phylogenetic (Morelli *et al*. 2016; Sol *et al*. 2017) and functional diversity (Lizée *et al*. 2011; Evans *et al*. 2018; Palacio *et al*. 2018), resulting in more homogenized wildlife communities. Despite these overall losses, cities can still harbor substantial biodiversity (Spotswood *et al*. 2021), including threatened species (Ives *et al*. 2016), with several factors contributing to an increase in species richness within urban areas. For example, biodiversity can be bolstered by green space (Beninde *et al*. 2015; Callaghan *et al*. 2018; Fidino *et al*. 2021), greater habitat heterogeneity (Oliveira Hagen *et al*. 2017; Souza *et al*. 2019), higher tree cover (Threlfall *et al*. 2016; De Castro Pena *et al*. 2017; Planillo *et al*. 2021), or reduced housing density (Fontana *et al*. 2011; Fidino *et al*. 2021). Within these species pools, some species – often termed urban adapters, urban exploiters, or urban-tolerant species – generally succeed in cities where others do not (Spotswood *et al*. 2021). The relative tolerance of species to urbanization can result from shared evolutionary history (Iglesias-Carrasco *et al*. 2022) and is often linked to functional traits. For example, in Australian birds, urban adapters show diet generalization, bigger brains and larger clutch sizes (Callaghan *et al*. 2019). Although many such traits have been suggested or regionally evaluated, what remains untested is whether the traits that confer urban tolerance in species differ across the cities and biogeographic contexts of the world. With recently-available global data on occurrence (Sullivan *et al*. 2009) and species trait (e.g., AVONET, Tobias *et al*. 2022), birds are an ideal system to explore this question.

Several ecological traits have been linked with urban association in birds (McClure 1989; Sol *et al*. 2014; Callaghan *et al*. 2019). For example, urban tolerance is often positively associated with niche breadth (Bonier *et al*. 2007; Evans *et al*. 2011), including dietary (Croci *et al*. 2008; Lizée *et al*. 2011; Morelli *et al*. 2016) and habitat breadth (Ducatez *et al*. 2018; Callaghan *et al*. 2019; Sayol *et al*. 2020). The degree of sociality also plays a role, with urban-tolerant species tending to be more social (Kark *et al*. 2007; Croci *et al*. 2008; Sol *et al*. 2014). In addition, nest placement is important, with ground nesters often avoiding urban areas (Conole & Kirkpatrick 2011; Evans *et al*. 2011; Sol *et al*. 2014; Dale *et al*. 2015; Guetté *et al*. 2017) while tree nesters tend to persist in cities (Conole & Kirkpatrick 2011; Dale *et al*. 2015). Yet, despite some general trends, the importance of certain traits often varies between studies. For example, although urban-associated species tend to have larger clutch sizes (Møller 2009; Lizée *et al*. 2011; Callaghan *et al*. 2019), this pattern is not always supported (Croci *et al*. 2008; Chamberlain *et al*. 2009), and may be mediated by other life-history traits (Sayol *et al*. 2020). Similarly, the role of body size has also received mixed support, with urban tolerance positively associated with body mass in Australia (Callaghan *et al*. 2019), but showing no relationship globally (Sol *et al*. 2017). Longevity or lifespan has seldom received strong support in models (Croci *et al*. 2008; Guetté *et al*. 2017), while cavity nesters show mixed responses to urban areas (Conole & Kirkpatrick 2011; Lizée *et al*. 2011; Dale *et al*. 2015; Jokimäki *et al*. 2016; Evans *et al*. 2018). Finally, although dispersal ability has been linked to urban tolerance (Møller 2009), migratory strategy is rarely associated with urban tolerance (Evans *et al*. 2011, 2018; Dale *et al*. 2015; Jokimäki *et al*. 2016; Guetté *et al*. 2017; Callaghan *et al*. 2019; Sayol *et al*. 2020).

The lack of generality in previous work may arise for multiple reasons. Many studies sample only a subset of biogeographic regions and/or species. Variation in the importance of traits may be driven by differences in species pools or by context-dependent differences in filters between different landscapes (Aronson *et al*. 2016; Oliveira Hagen *et al*. 2017). It thus seems likely that results should differ between biomes due to differences in climate and biogeographic history (Morelli *et al*. 2016; Leveau *et al*. 2017; Filloy *et al*. 2019). Yet, even studies that have taken a global perspective have been biased in their sampling towards North America, Europe, and Australia, with a distinct lack of data from the tropics (Magle *et al*. 2012; Sol *et al*. 2014; Sayol *et al*. 2020). Moreover, the number of species in global trait studies has also been limited, with the largest sample size (629 species by Sayol et al. 2020)) representing only ∼6% of bird species found globally. Previous studies have been restricted by the lack of bird occurrence data across urbanization gradients, particularly in the tropics (Magle *et al*. 2012), but also by access to global trait datasets that have only recently become available.

Here, we combine global data on occurrence (>125 million records) from the citizen science project eBird (Sullivan *et al*. 2009) with a continuous measure of urbanization (night-time lights) to calculate an Urban Association Index (UAI) for 3768 bird species (∼35% of extant bird species) in 137 cities across six continents and 11 biomes. We chose ten species-specific functional traits with globally available data and hypothesized links to urban tolerance, and modeled UAI values as a function of these traits. We further chose three city-specific landscape variables that we predicted would influence the importance of our traits for urban tolerance (Oliveira Hagen *et al*. 2017), assessing whether the effects of each trait varied as a function of latitude, human population density, and landscape greenness. We present the first evidence that the importance of different traits for urban tolerance varies predictably across the planet.

## Methods

### Data filtering

We downloaded the global eBird basic dataset (Sullivan *et al*. 2009) including all records up until February, 2022 (v1.14). We restricted the dataset to the years 2002–2021 – the 20 complete years before present. We then limited eBird protocol types to “traveling”, “stationary”, and “area” and removed incomplete checklists. Following eBird best practices (Johnston *et al*. 2021), we removed checklists with >10 observers, with durations >5 hr, with distances >5 km (for “traveling” protocol), and with areas >500 ha (for “area” protocol). For group checklists involving duplicate records, we randomly retained one checklist per group. Finally, we removed records that were not identified to species level, including all hybrids, intergrades, “slashes” (e.g., “Greater/Lesser Yellowlegs”), indefinite species (e.g., “hummingbird sp.”), and domestics. Although many of the species in our dataset are introduced in some cities, they are native in others, so we did not remove or classify species based on being exotic (e.g., *Passer domesticus, Sturnus vulgaris*). We made a single exception to these exclusions, retaining the widespread and ubiquitous Feral (Rock) Pigeon (*Columba livia*) despite having been domesticated as it is a key avian species in many cities. Initially we considered including all species found in cities but restricted our dataset to exclude water birds (∼15% of the species set) since they have substantially different natural histories and traits compared to land birds, following Callaghan et al. (2019).

### City selection

From the data repository OpenDataSoft, we downloaded the dataset “Geonames – All Cities with a population > 1000” (OpenDataSoft 2022), and reduced the dataset down to cities with a population >100,000, yielding 4643 cities. We chose this relatively low population cut-off to include smaller, remote cities in ecologically distinct regions – including Darwin (Australia), Port Louis (Mauritius), and Reykjavík (Iceland). We then calculated the pairwise distance between every city using the package *geodist* (Padgham & Sumner 2019). Starting with the cities with the largest populations, we sequentially removed all smaller cities within 500 km of the larger city in order to produce a set of non-overlapping, spatially-independent cities. This algorithm retained 289 cities separated by at least 500 km (Fig. S1). After identification of these target urban areas around the world, we filtered the eBird dataset to checklists within a 100 km radius of each city center. This radius was chosen to include the whole metropolitan area as well as surrounding habitats that might supplement the species pool. For each city dataset (hereafter “city”), we removed species with <100 records, as well as species that comprised <0.01% of all occurrences per city. The first filter ensured a minimum data requirement while the second filter was a threshold intended to filter out vagrant species while retaining scarce but expected species. As some cities lacked 100 records for even one species, we removed any city with <50 species remaining after restricting species to ≥100 records, such that all remaining cities had ≥5000 bird records. This 50-species threshold was chosen in order to remove cities that contained only a handful of species that would tend to be more urban associated (high UAIs), but to retain cities in environments with low species richness (e.g., boreal regions) that would have been removed if the threshold was 100 species. Our final dataset contained 16,455 UAI estimates representing 127,046,578 eBird occurrence records of 3768 species across 137 cities (Fig. 1a).

**Figure 1.**
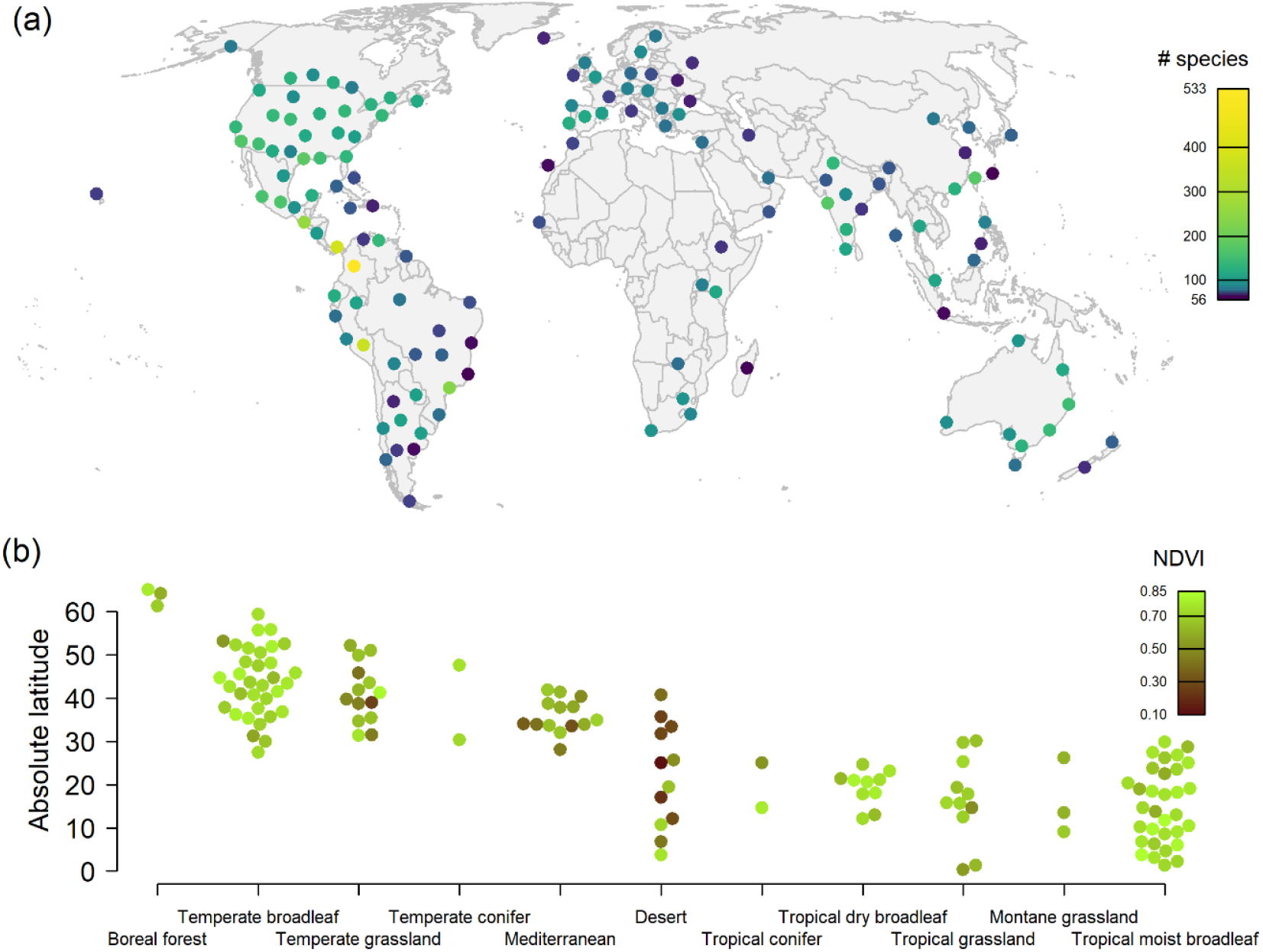
Geographical coverage of 137 cities included in the analysis. Cities were distributed (a) across 62 countries, including 39 in North America, 28 in South America, 27 in Asia, 22 in Europe, 10 in Africa, and 10 in Australasia. Each of these cities was initially selected to have a population of ≥100,000 people and be ≥500 km apart. Cities were then retained that had ≥50 species each with ≥100 eBird records within a 100 km radius circles over 20 years (2002–2021). Cities are colored by the (log) number of species that met the criteria from 56 (dark purple) to 533 (yellow). Cities were representative (b) of 11 of the world’s biomes (Olson & Dinerstein 1998). Biomes are ordered by the mean absolute latitude of the cities included, and cities are colored by the NDVI of the greenest month, from the greenest city (lime green) to the least green city (dark brown).

### Urbanization association index (UAI)

To quantify species’ relationships with urban areas we aimed to create a continuous metric of urban association that would avoid the assumptions of using arbitrary thresholds to categorize species based on urban tolerance (Callaghan *et al*. 2019; Fanelli *et al*. 2022). Following Callaghan et al. (2019), we downloaded the VIIRS night-time lights imagery (Annual VNL V1; Elvidge et al. 2017), a composite global image of night-time lights for the year 2016. Although 2016 is not the mid-point of the eBird data (2012), the number of eBird records has increased exponentially so 2016 is close to the mean year of all checklists (2017). We chose light radiance as a proxy for urbanization because it is available as a continuous measure across the world, it is a close proxy for human population density (Elvidge *et al*. 2017), and, when combined with eBird data, it is correlated with other measures of urban tolerance in birds (Callaghan *et al*. 2021). From this imagery, we extracted the radiance value for every eBird checklist locality. As these radiance values start at 0 (total darkness) and increase exponentially, we added 1 then log-transformed all radiance values to reduce the leverage of extremely bright buildings. Then, for every species within each city, we calculated the mean radiance value of all occurrence records. We chose to use the mean instead of the median because we found that many species had a median radiance of 0, as they occurred predominantly in non-urban areas. Moreover, the distribution of mean radiance values was fairly normal (Fig. S2), while the distribution of median radiance values was heavily right skewed. We also tested our chosen metric for sensitivity to mismatches in scale between the spatial resolution of VIIRS and checklists (Supporting Information) but decided to retain the metric as described. Thus, our Urban Association Index (UAI) for each species is the mean of the transformed radiance values across eBird records where the radiance value of each record is taken from a single pixel of night-time lights.

### Species traits

We chose species-specific functional traits that have been linked to urban tolerance in the past and/or traits that we hypothesized would predict urban tolerance that have not been tested globally. We chose traits that were available for the entire species set and, where possible, we chose numerical (rather than categorical) traits in order to reduce the number of parameters estimated. We therefore did not use traits such as residual brain size where data does not exist for all species (Sayol *et al*. 2020) and excluded categorical traits with many levels, such as primary diet. Traits for every species were then extracted from several datasets, as follows (Table 1).

**Table 1.** Ten functional traits included in the analysis.

| Trait | Description | Primary sources |
| --- | --- | --- |
| Body mass | Log-transformed | AVONET |
| Bill shape | Second PC from a PCA of four bill measurements | AVONET |
| Hand-wing index | The ratio of Kipp's distance to wing length | AVONET |
| Diet breadth | Number of major food groups, 1–9 | BirdBase, Birds of the World |
| Habitat breadth | Number of major habitats, 1–11 | BirdBase, Birds of the World |
| Lower elevational limit | Lower limit of elevational range reported in the literature | BirdBase, Birds of the World |
| Territoriality | A scale from 1 (low) to 3 (high) | Tobias et al. 2016 |
| Longevity | Log-transformed | Bird et al. 2020 |
| Clutch size | Log-transformed | BirdBase, Myhrvold et al. 2015, Birds of the World |
| Nest type | Categorical: ground, cavity, open, and enclosed | BirdBase, Birds of the World |

From the publicly available AVONET (Tobias *et al*. 2022) we extracted body mass, four bill measurements (length from culmen, length from nares, width, and depth), and hand-wing index (HWI). These data were complete for all species. To reduce the four bill measurements down to a single axis, we conducted a PCA on the variables and extracted the second principal component, ignoring the first principal component, which is highly correlated with body size (Pigot *et al*. 2020). This second principal component – which we refer to as “bill shape” – represents a spectrum from long, thin, pointy bills (e.g., *Ensifera ensifera*) to short, thick bills (e.g., *Callocephalon fimbriatum*), a spectrum associated with foraging specializations (Pigot *et al*. 2020). As a measure of dispersal ability, HWI has not been tested as a global predictor of urban tolerance but is highly correlated with several ecological factors, including primary diet and habitat type (Sheard *et al*. 2020).

From the dataset BirdBase (Şekercioğlu *et al*. 2004; Buechley *et al*. 2019) we extracted diet breadth, habitat breadth, lower elevational limit, clutch size, and nest shape/substrate. Diet breadth is the number of major food groups (1–9) that a species eats (e.g., invertebrates, fruit, seeds) while habitat breadth (1–11) is the number of major habitats where a species is found (e.g., forest, grasslands, desert). Lower elevational limit was included because we hypothesized that cities – which tend to be found non-randomly at lower elevations (Luck 2007) – would favor species that occur at lower elevations. Nest shape and nest substrate were originally sourced as two separate data columns, but we collapsed these into one. As there was no way to define these nests numerically by shape and substrate, we instead defined four categories: ground (nests of any form located on the ground), cavity (nests above ground in cavities or crevices), open (nests above ground with open tops such as cups, saucers, and platforms), and enclosed (nests above ground with entrance holes such as spheres, pendants, and domes). Clutch size data were augmented with data from an existing published dataset (Myhrvold *et al*. 2015), while further gaps in BirdBase variables were filled using the online database Birds of the World (Billerman *et al*. 2020). Where information was lacking for a species, missing values were inferred from close extant relatives with complete data. Finally, longevity (a measure of lifespan) and territoriality (a scale from 1 to 3 where 3 is more territorial) were extracted from published datasets (Tobias *et al*. 2016; Bird *et al*. 2020). Once assembled, we had complete data for ten functional traits (Table 1).

Trait variables were transformed, as necessary, prior to analysis. Given expected non-linear relationships, we took the log of body mass, longevity, and clutch size. We then scaled and centered all numerical traits to have a mean of 0 and a standard deviation of 1.

### City variables

For each 100 km-radius city circle we gathered data on three covariates that we hypothesized would alter the importance of traits: latitude, greenness, and population density. We chose numerical covariates in order to reduce the number of parameters, as each new city covariate adds nine parameters (one for each numerical trait) to the model. However, combined, latitude and greenness cover much of the variation among biomes (Fig. 1b).

Many factors vary with latitude including climate, species richness, and human development, so there are many possible avenues through which latitude could affect urban tolerance. For example, the stability of tropical climate and ecosystems (Janzen 1967) may mean stronger filters in urban areas against ecological specialists in the tropics compared to temperate regions (Newbold *et al*. 2013). We extracted the latitude of each city from the same Geonames dataset as the city populations.

The amount of greenness in a city – whether tree cover or vegetation diversity – is an important predictor of bird diversity in cities (Beninde *et al*. 2015; Threlfall *et al*. 2016; De Castro Pena *et al*. 2017; Callaghan *et al*. 2018; Souza *et al*. 2019; Planillo *et al*. 2021). Moreover, overall greenness of the landscape depends on the primary habitat. For example, desert cities such as Phoenix (USA) and Dubai (UAE) are greener than the surrounding landscape while forest cities such as Iquitos (Peru) and Nashville (USA) are less green than the surroundings. We thus hypothesized that the amount of greenery would also alter trait filters (Oliveira Hagen *et al*. 2017). For example, less green landscapes with fewer resources may select for habitat generalists or more mobile species. We used NDVI as a measure of the greenness of each city, derived from the MOD13A3 product (Didan 2021). This product provides 1km monthly NDVI values globally, excluding water bodies. We calculated the mean NDVI values within the 100 km buffer of each city for each month for the year 2021, and retained the maximum NDVI value. We used the maximum NDVI value as each city has a different seasonal cycle over which greenness is likely to vary (i.e., greenness peaks in some cities in August, while in January in others).

Human population density has been linked to taxonomic and functional diversity in cities (Fontana *et al*. 2011; Oliveira Hagen *et al*. 2017). We hypothesized that cities with higher population densities may present strong selection pressures against species that are, for example, larger with narrower diets. To obtain population density (number of people/cell), we downloaded Gridded Population of the World data from the Center for International Earth Science Information Network (CIESIN 2018). The data are available on 5-year intervals between 2000–2020. We used 30 arc-second resolution population size for the year 2015 as the year closest to the VIIRS imagery and the mean year of eBird records. We buffered city midpoints by 100 km and extracted the mean value of the gridded density data within each buffer.

For the models, we calculated the absolute value of latitude and the log of population density. All three city covariates were then scaled and centered.

### Modeling

We modeled UAI values as a function of traits and city variables in a Bayesian hierarchical framework that accounted for the random effects of city and species. We modeled the effect of the ten species traits on UAI with the following structure:

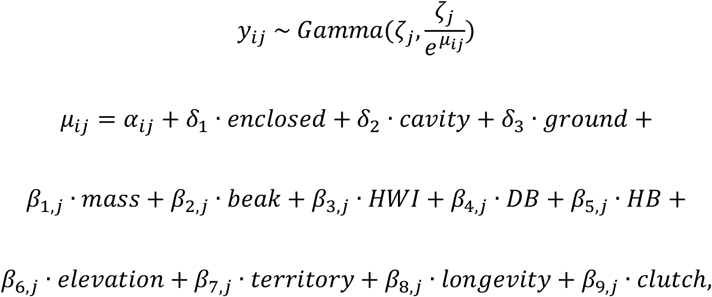

where the estimated mean UAI, *y*_*ij*_, for species *i* in city *j* was modeled as a gamma-distributed random variable with a city-specific shape parameter *ζ*_*j*_ and a rate parameter equal to 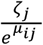. We chose a gamma distribution to reflect the fact that our response variable was bounded by 0 on the lower end and right skewed. The shape of the distribution was allowed to vary among cities to accommodate variation in the data. In turn, *μ*_*ij*_ was modeled as a linear combination of an intercept for open nesters, *α*_*ij*_, three differences in intercepts (*δ*_1_ to *δ*_3_) and nine covariates with corresponding parameters (*β*_1,*j*_ to *β*_9,*j*_). The parameters *δ*_1_ to *δ*_3_ represent the difference in UAI for three dummy variables (*enclosed, cavity*, and *ground*) that together encode the three other nest types, where all three covariates are binary (1 = species’ nest type, 0 = otherwise) and mutually exclusive. The parameters *β*_1,*i*_ to *β*_9,*i*_ represent the slopes of the effects of nine numerical traits on *μ*_*i*_.

The intercept *α*_*i*_ can be further decomposed,

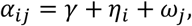

into a global intercept, *γ*, and the random effects of species, *η*_*i*_, and city, *ω*_*j*_. The random effect of species accounts for species being represented across multiple cities. The random effect of city allows species in different cities to have different average UAIs based on unmodeled factors such as differences in brightness, development, and birdwatching effort. The random effects of species,

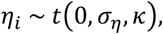

were drawn from a t-distribution with a mean 0, standard deviation *σ*_*η*_, and degrees of freedom *κ* (which controls the degree to which the distribution resembles a normal, as *κ* approaches infinity, or a Cauchy, as *κ* approaches 1). The choice of t-distribution allowed for fatter tails in the distribution of intercepts across species. The random effects of city,

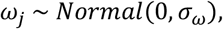

were drawn from normal distributions with mean 0 and standard deviation and *σ*_*ω*_.

In order to integrate the city covariates with the functional traits, every *β*_*j*_ parameter for trait *k* was drawn from a normal distribution

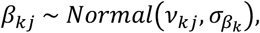

with a mean *v*_*kj*_ and a standard deviation 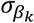 such that each *β*_*ki*_ had its own process error, to accommodate variation in the data. The mean was then modeled as a linear combination of city covariates

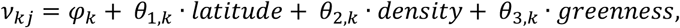

such that the effect of each numerical trait on UAI varied as a function of the city-level covariates. Importantly, this allowed urban tolerance to be predicted differently by different traits in different geographical contexts.

We ran this Bayesian model using the program JAGS (Plummer 2003) via the R package *R2jags* (Su & Yajima 2021). We used vague priors (mean of 0, standard deviation of 100) and we ran three chains, each with 40,000 iterations, beginning with a burn-in of 10,000 followed by a thinning of 30, retaining 1000 posteriors per chain. We verified that the model had successfully converged (Rhat ≤ 1.01 and n.eff > 400). We performed posterior predictive checks to ensure that data generated by the model were similar to data used to fit the model (Gelman *et al*. 2000). We found that 49.4% of the means of the posteriors were less than the observed mean UAI, indicating that our model could successfully reproduce the mean UAI. From the parameter posteriors we calculated the means and 95% credible intervals for each parameter. Due to Bayesian shrinkage within the random effects framework, *post hoc* testing indicated that the model was less able to estimate suitable species-specific intercepts, *η*_*i*_, for species with only a few data points – i.e., those species present in only one or a few cities. We therefore fit a second, identically-structured model using a dataset where species represented in <5 cities were excluded, and checked for consistency of model results (i.e., *β*s) as compared to the original, full dataset model.

### Phylogenetic structure

In order to test for phylogenetic signal in UAI values, we aligned the eBird (Clements) taxonomy with the taxonomy of BirdTree.org (Jetz *et al*. 2012) and downloaded 100 phylogenetic trees with the Hackett backbone. We averaged UAI values across the tips of the phylogeny. For each tree, we calculated Pagel’s *λ* as a measure of phylogenetic signal using the package *phytools* (Revell 2012). We then calculated the mean *λ* across trees, with associated 95% quantiles. Directly incorporating these phylogenies into our Bayesian model was not practical due to the extensive computational time required for an analysis that includes so many species (over a year). Rather, we tested whether model residuals, averaged at the species level, contained phylogenetic signal (Revell 2010). This test would tell us whether there was unexplained variation in the model associated with phylogeny. We also tested for signal in the residuals of the model with the reduced species set.

## Results

Our analysis included 16,455 UAI estimates representing data from >125 million eBird records across 137 cities (Fig. 1a). This list comprised cities from 62 countries including 39 in North America, 28 in South America, 27 in Asia, 22 in Europe, 10 in Africa, and 10 in Australasia. Together, these cities span 11 of the world’s 14 terrestrial biomes (Olson & Dinerstein 1998). The number of avian species meeting the inclusion criteria in each city ranged from 56 in Naha (Japan) to 533 in Bogotá (Colombia).

Of the 3768 species for which we calculated UAI, the five species present in the most urban areas (see Fig. S4 for top 30) were Feral (Rock) Pigeon (*Columba livia*), House Sparrow (*Passer domesticus*), Barn Swallow (*Hirundo rustica*), Osprey (*Pandion haliaetus*), and Peregrine Falcon (*Falco peregrinus*). Across species, UAI values ranged from 0 (for 46 species) to 3.97 (Yellow-crested Cockatoo – *Cacatua sulphurea* – a species introduced to Hong Kong) with a mean of 1.14. Of species present in at least ten cities, the top-five species with the highest UAI (see Fig. S5 for top 30) were Monk Parakeet (*Myiopsitta monachus*), Rose-ringed Parakeet (*Psittacula krameri*), Yellow-chevroned Parakeet (*Brotogeris chiriri*), Feral Pigeon, and Sayaca Tanager (*Thraupis sayaca*).

There was considerable phylogenetic signal in UAI across species (Fig. 2; *λ* = 0.61, CI = 0.56–0.65). Notable families with high average UAI values, indicating broad urban associations, included Sturnidae (starlings; 1.75 ± 0.13 SE, n_cities_ = 40), Apodidae (swifts; 1.61 ± 0.12, n_cities_ = 44), Hirundinidae (swallows; 1.55 ± 0.09, n_cities_ = 52), Psittacidae (parrots; 1.55 ± 0.11, n_cities_ = 86), and Icteridae (New World orioles and blackbirds; 1.47 ± 0.08, n_cities_ = 80). Notable families with low average UAI included Pipridae (manakins; 0.33 ± 0.07, n_cities_ = 21), Petroicidae (Australasian robins; 0.38 ± 0.09, n_cities_ = 20), Trogonidae (trogons; 0.45 ± 0.07, n_cities_ = 24), Thamnophilidae (antbirds; 0.55 ± 0.06, n_cities_ = 72), and Tinamidae (tinamous; 0.58 ± 0.09, n_cities_ = 22).

**Figure 2.**
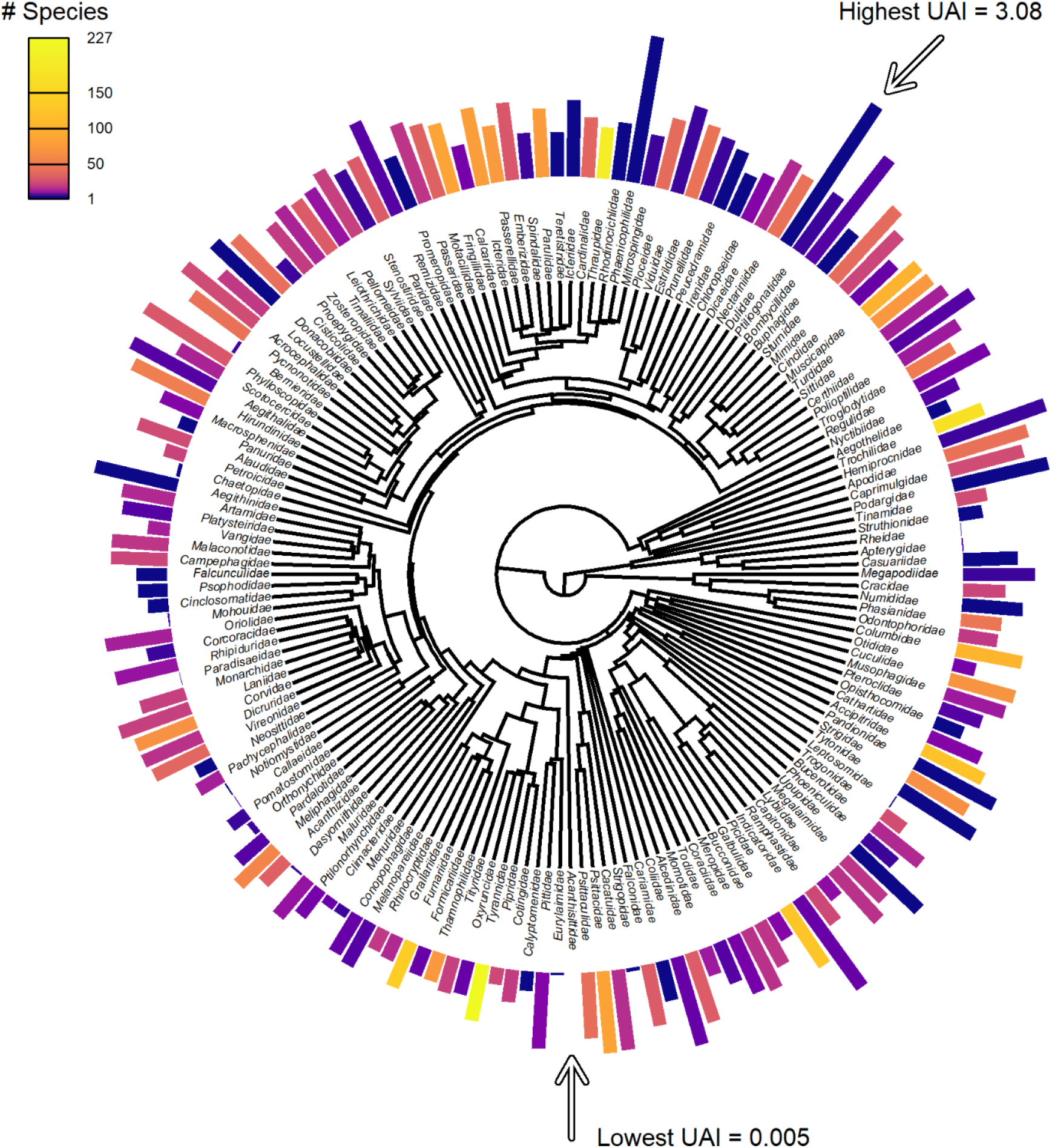
The phylogeny of Urban Association Index (UAI) across 3768 species by family. For visualization, UAI values were averaged across species and then across taxonomic families. The height of the bar indicates the UAI, with taller bars indicating higher urban tolerance. The color indicates the (log) number of species in the family from 1 (dark purple) to 231 (Tyrannidae, yellow).

Of the ten species-specific traits considered, all except bill shape were significantly associated with UAI (Figs. 3,4). Body mass (Fig. 3a), lower elevational limit (Fig. 3e), territoriality (Fig. 3f), and ground nesting (Fig. 3i) were negatively associated with UAI, while hand-wing index (HWI; Fig. 3b), diet breadth (Fig. 3c), habitat breadth (Fig. 3d), longevity (Fig. 3g), and clutch size (Fig. 3h) were positively associated with UAI. In other words, more urban-tolerant species are smaller, tree- or building-nesting species with higher dispersal ability, wider diet and habitat breadth, lower elevational limits, lower territoriality, longer lifespan, and greater clutch size.

**Figure 3.**
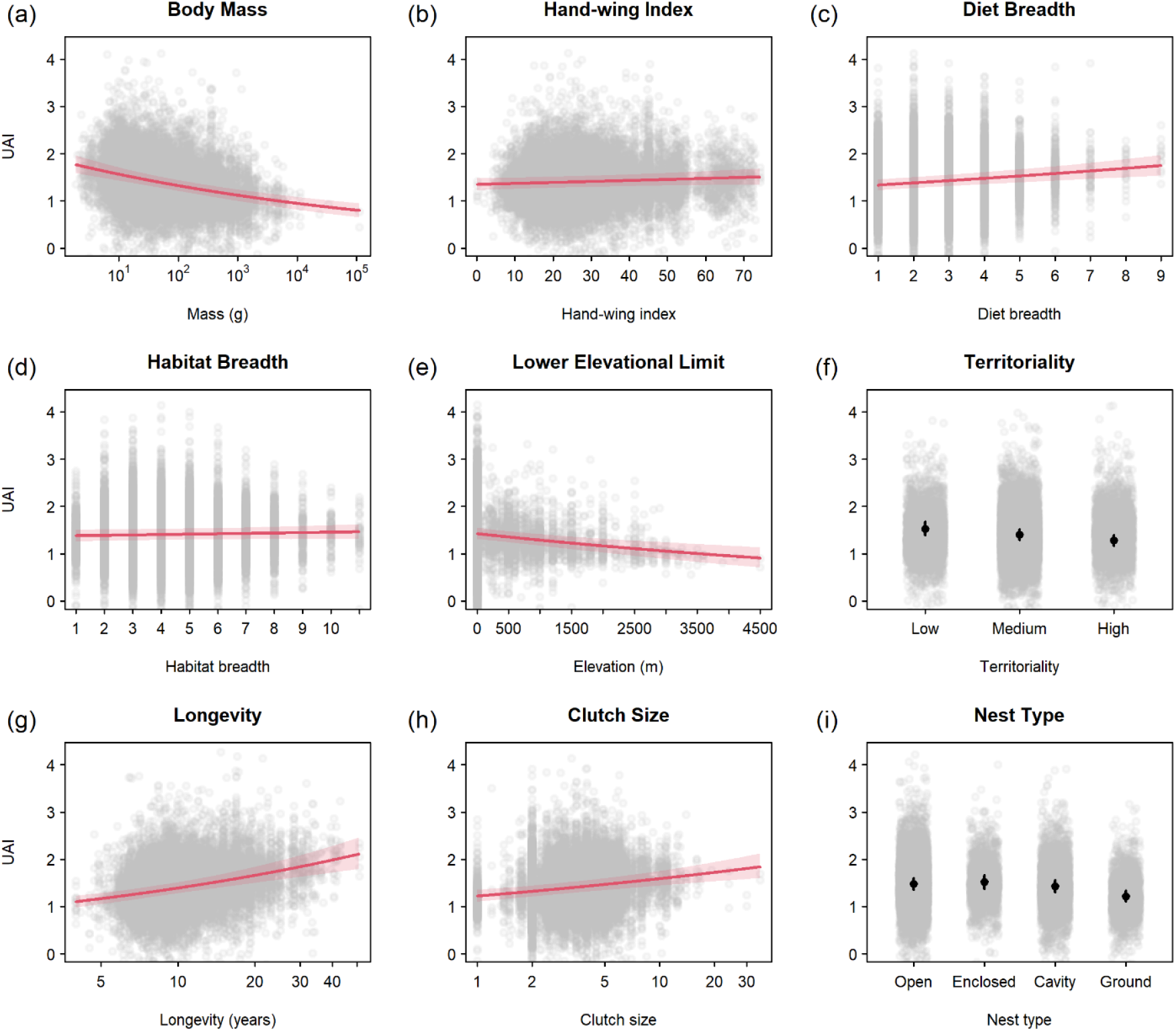
The global mean effects of nine species traits on the Urban Association Index (UAI) of 3768 bird species across 137 cities. There was a significant relationship between UAI and (a) body mass, (b) hand-wing index, (c) diet breadth, (d) habitat breadth, (e) lower elevational limit, (f) territoriality, (g) longevity, (h) clutch size, and (i) nest type. Gray points show the partial residuals of each data point. Trend lines for numerical traits are shown along with the 95% credible intervals. Territoriality is treated as numerical in the model but here we summarize the data for the three levels of territoriality (low, medium, high). Nest type is treated as categorical in the model (open, enclosed, cavity, or ground). For territoriality and nest type, black points show the mean and bars show the 95% credible intervals.

**Figure 4.**
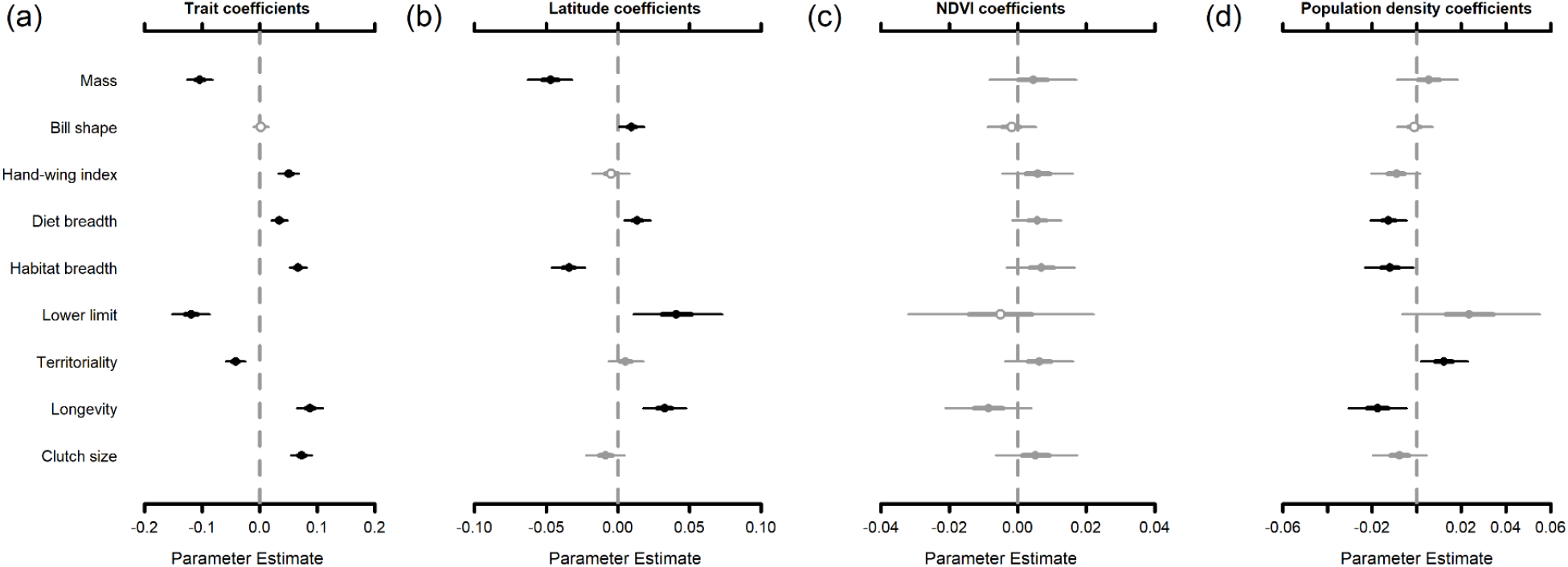
The relationship between Urban Association Index (UAI), species traits, and city variables for 3768 birds species across 137 cities. Covariate coefficients (*β*) show how UAI varies as a function of nine numerical trait covariates (a). In turn, these trait coefficients vary across cities as a function of three city-level variables with corresponding coefficients (*θ*): (b) latitude, (c) NDVI, and (d) human population density. Points show the covariate coefficient estimates with corresponding interquartile range (thick lines) and 95% credible intervals (thin lines). Points are open when the interquartile range overlaps 0. Points and lines are gray when the 95% credible intervals overlap 0 and black when they do not.

Seven of the traits varied significantly as a function of city-level covariates (Fig. 4,5). In terms of latitude (Fig. 4b), the negative effect of body mass on UAI (Fig. 5a) and the positive effects of diet breadth (Fig. 5b) and longevity (Fig. 5d) became more pronounced in cities at higher latitudes. Contrastingly, the positive effect of habitat breadth (Fig. 5e) and the negative effect of lower elevational range limit (Fig. 5g) on UAI became more pronounced in tropical cities. The effect of bill shape on UAI – which showed no globally consistent relationship – varied with latitude (Fig. 5h) such that species with longer, pointier bills were more urban tolerant at higher latitudes while species with shorter, thicker bills were more urban tolerant at lower latitudes. In terms of population density (Fig. 4d), the negative effect of territoriality (Fig. 5c) and the positive effects of diet breadth (Fig. 5f) and longevity (Fig. 5i) became more pronounced in cities with lower population density. Finally, none of the nine numerical traits varied significantly in effect as a function of landscape greenness (Fig. 4c).

**Figure 5.**
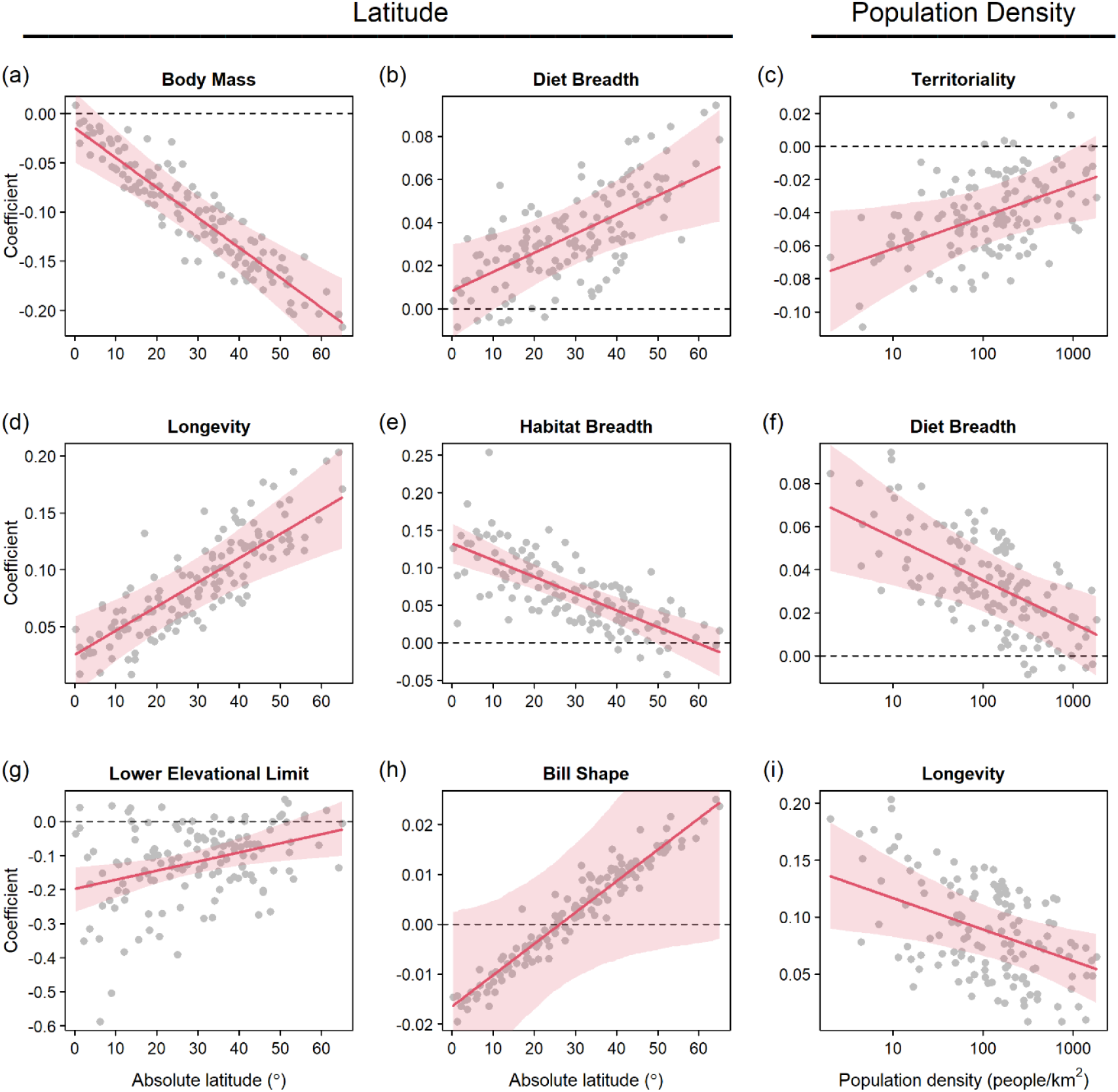
The effect of latitude and human population density on the city-level trait coefficients for Urban Association Index. Latitude had a significant effect on the city-level trait coefficients for (a) body mass, (b) diet breadth, (d) longevity, (e) habitat breadth, (g) lower elevational limit, and (h) bill shape. Human population density had a significant effect on the city-level trait coefficients for (c) territoriality, (f) diet breadth, and (i) longevity. Points represent the model-estimated trait coefficients for each city (n = 137). Trend lines and 95% credible intervals show how these coefficients vary as a function of the city-level covariates.

Model residuals contained relatively low phylogenetic signal (*λ* = 0.37, CI = 0.33–0.43). Most of this signal resulted from species with few data points – i.e., those represented in 1–4 cities – as the model was less able to estimate suitable species-specific intercepts due to the shrinkage of intercept parameters towards the cross-species mean. Removing these 2848 species (76% of the species set) and re-running the model produced qualitatively similar trait coefficients (Fig. S6) and greatly reduced the phylogenetic signal in the residuals (*λ* = 0.15, CI = 0.09–0.22). Thus, we are confident that our estimates of the effect of traits on UAI are robust to potential phylogenetic or sample-based biases.

## Discussion

Many studies have linked species-specific functional traits to urban tolerance (Møller 2009; Sol *et al*. 2014; Callaghan *et al*. 2019; Sayol *et al*. 2020) but none have tested for interactions between traits and geographic factors, especially not at the global taxonomic and spatial scale we employ here. For 35% of the world’s bird species across 137 cities and 11 biomes – including regions of the world underrepresented in ecological studies (i.e., Asia, Africa, South America; Magle et al. 2012; Estes et al. 2018) – we find that nine different functional traits are related to urban association. Furthermore, we find that two geographic variables – latitude and human population density – significantly modulate the effects of seven of these traits, meaning that the strength of trait-based filters in urban environments varies systematically across the planet (Aronson *et al*. 2016; Oliveira Hagen *et al*. 2017). Our study is the first at a global scale to demonstrate the effects of body size, hand-wing index (HWI), diet breadth, lower elevational limit, territoriality, longevity, and clutch size on urban association, and confirms the positive association of habitat breadth and the negative association of ground nesting (Sol *et al*. 2014; Ducatez *et al*. 2018).

Urban associated species tended to have wider diet and habitat breadths (Fig. 3c,d), confirming the role of ecological generalism in urban tolerance (Bonier *et al*. 2007; Ducatez *et al*. 2018; Callaghan *et al*. 2019; Fidino *et al*. 2022). As cities erase or erode most native habitats (McDonald *et al*. 2020), ecological specialists are less able to survive while more versatile species persist. However, we found that the importance of diet and habitat breadth had opposing patterns across latitude (Fig. 5b,d). Habitat breadth was more important in tropical urban areas, possibly because most tropical land birds have high forest dependency (Tobias *et al*. 2013), and thus are more likely to experience a stronger filter in urban areas (Newbold *et al*. 2013). But, with fewer habitats to specialize on towards the poles, habitat breadth becomes less important at higher latitudes. By contrast, diet breadth was more important in temperate areas. Many urban-associated tropical birds are dietary specialists, particularly nectarivores and frugivores, where they take advantage of plentiful year-round fruiting and flowering ornamental trees (Lim & Sodhi 2004). Temperate cities, with seasonal resource pulses and troughs, favor omnivores that can make use of a wide variety of food sources (Croci *et al*. 2008; Lizée *et al*. 2011; Jokimäki *et al*. 2016; Evans *et al*. 2018).

Related to diet, the effect of beak shape on urban associations changed sign with latitude (Fig. 5f). In the tropics, species with short, thick bills were favored in urban areas, a result that may be explained by the abundance of specialist frugivores in fruit-plentiful tropical cities, exemplified by urban-tolerant parrots (Cacatuidae, Psittaculidae, Psittacidae). At more temperate latitudes, species with short, stubby bills tend to be granivores and also tend to avoid urban areas where grasses are cut short. While the occassional short, stubby bill does well in temperate urban environments (e.g., House Sparrow or House Finch, *Haemorhous mexicanus*), many temperate granivores such as game birds (Phasianidae), longspurs (Calcariidae), and grassland sparrows (Passerellidae) require suitable habitat far from development (Croci *et al*. 2008; Callaghan *et al*. 2019). Meanwhile, the hummingbirds (Trochilidae) – long-billed species with high data leverage – present an interesting outlier. In the Neotropics, where their diversity peaks, only a fraction of species are found in urban areas (Maruyama *et al*. 2019) such as Panama City, while in North America, most hummingbird species frequent urban feeders (Greig *et al*. 2017; Miller *et al*. 2017). Variation in the importance of bill shape is clearly complex, underscoring the diverse responses of different feeding guilds to urbanization (Kark *et al*. 2007; Jokimäki *et al*. 2016; Evans *et al*. 2018; Callaghan *et al*. 2019).

Previous studies have suggested that migratory strategy was not associated with urban tolerance (Dale *et al*. 2015; Jokimäki *et al*. 2016; Guetté *et al*. 2017; Evans *et al*. 2018; Callaghan *et al*. 2019; Sayol *et al*. 2020), while only one study has found dispersal ability *per se* to be related to urban associations (Møller 2009). We, however, found that species with higher HWI, i.e., longer, more pointed wings associated with greater dispersal ability (Sheard *et al*. 2020), have higher UAI values (Fig. 3b). Although dispersal ability is positively associated with migratory capacity (Sheard *et al*. 2020), previous studies focusing on temperate cities may not have found a role for migratory capacity as migrants tend to broaden their habitat use to include cities on their wintering grounds in the tropics. Additionally, this pattern could be driven by a number of factors, including the sensitivity of low-dispersal species to anthropogenic change (Claramunt *et al*. 2022), and the association between HWI and specific foraging modes such as flycatching, aerial insectivory, frugivory, and nectarivory (as opposed to gleaning, terrestrial insectivory etc.) that would be favored in urban environments (Lees & Peres 2009; Sheard *et al*. 2020).

The role of body size in urban tolerance has mixed support, including a positive association in Australia (Callaghan *et al*. 2019) but no effect globally across 358 species (Sol *et al*. 2014). Here, we found urban-tolerant species are significantly smaller, an effect (Fig. 3a) that strengthens towards the poles (Fig. 5a). Many families of large species, such as bustards (Otididae), tinamous, and pheasants, appear to be urban avoidant (Fig 2). These species tend also to be cursorial, which could put them at elevated risk of urban-associated predators (e.g., domestic cats; Loss et al. 2013) and nest predators (e.g., rats; Smith et al. 2016). In the tropics, these families of large species might be balanced out by urban-tolerant arboreal-nesting large hornbills (Bucerotidae), turacos (Musophagidae), parrots, and cockatoos (Conole & Kirkpatrick 2011). In temperate regions, game birds are likely selected against in urban areas due to habitat requirements, the history of hunting, or pressure from meso-predators (Crooks & Soule 1999).

Body size can be associated with other life-history traits that predict urban tolerance – although we found little correlation between body mass, longevity, and clutch size in this study. Supporting results from other studies (Møller 2009; Lizée *et al*. 2011; Callaghan *et al*. 2019), we found that species with larger clutches were more urban tolerant (Fig. 3h). Species with larger clutch sizes tend to live at the faster end of the life-history continuum and may be able to adapt faster to novel environments (Møller 2009). Conversely, however, we found that species with longer lifespans were also more urban tolerant (Fig. 3g), corroborating the finding that urban-tolerant species also have higher annual survival rates (Møller 2009). One possibility is that long-lived species are also more intelligent species (Smeele *et al*. 2022). The role of brain size in urban tolerance appears linked to other life-history strategies, with big brains important for species with high brood value (i.e., fewer broods over a lifetime) and small brains important for species with low brood value (Sayol *et al*. 2020). While we lacked the data to test this hypothesis globally across our full species set, our results suggest a similar trade-off, that it helps to either have large clutch sizes, or live longer in order to learn to exploit urban environments. The importance of longevity also increases in temperate cities (Fig. 5c), suggesting that living longer and, perhaps, being smarter are more beneficial closer to the poles.

Certain aspects of breeding biology were also tied to urban tolerance. While we did not test sociality *per se* (a trait which is not available broadly), we did find a significant negative effect of territoriality (Fig. 3f). Urban-tolerant species tend to be more social or gregarious (Kark et al. 2007; Croci et al. 2008; Sol et al. 2014) and therefore less territorial. Being strongly territorial year-round (level 3 on the scale) is usually tied to defense of resources (Tobias *et al*. 2016) and in resource-poor cities it makes less sense to be territorial and more sense to follow resources more plastically. Where species nest also matters, and we confirm the results of others that ground-nesting species tend to be less urban-tolerant (Conole & Kirkpatrick 2011; Evans *et al*. 2011; Sol *et al*. 2014; Dale *et al*. 2015; Guetté *et al*. 2017). Species that nest above the ground with open or enclosed nests had the highest urban tolerance (Fig. 3i), probably due to safety from predators (Jokimaki & Huhta 2000; Chace & Walsh 2006). Some studies have suggested that cavity nesters would have higher urban tolerance (Chace & Walsh 2006; Croci *et al*. 2008; Conole & Kirkpatrick 2011) while others have suggested the opposite (Evans *et al*. 2018). We found intermediate UAI values for cavity nesters, perhaps reflecting the contrast of relative success of cavity nesters with lower availability of nest cavities in urban areas (Blewett & Marzluff 2005).

The effects of territoriality, diet breadth, and longevity were all reduced in cities with higher population density (Fig. 5g–i). As population density is calculated across the whole 100 km radius circle, it is possible that the most densely populated cities are more homogenous with less non-urban habitat for urban avoiders. For example, Anchorage (USA) and Reykjavík (Iceland) are small cities surrounded by wilderness where habitats strongly differ between urban and non-urban areas. In contrast, cities like Bangkok (Thailand) and İstanbul (Turkey) are vast sprawling metropolises with abundant feral predators where there is little room for specialized, long-lived, territorial urban avoiders. Finally, we did not find that any trait effects varied as a function of landscape greenness, indicating that ecological filters in urban areas are similar for cities at the same latitude that differ in greenness, a measure driven in large part by habitat (Fig. 1b).

In summary, we found that numerous species-specific functional traits (smaller body size, lower territoriality, greater dispersal ability, broader dietary and habitat niches, larger clutch sizes, greater longevity) predict urban tolerance across the planet. However, many of these trait effects are modulated by landscape-level properties, most notably latitude. Where previous studies have demonstrated the importance of certain traits in certain parts of the world, we demonstrate the importance of geography in determining trait-based urbanization filters (Ferenc *et al*. 2014; Aronson *et al*. 2016; Leveau *et al*. 2017; Filloy *et al*. 2019) at an unprecedented taxonomic and spatial scale. Moreover, much of the region-specific variation in previous trait-seeking studies could be due to predictable geographic variation in trait strength that varies with latitude and human population density. Studying how traits filter diversity across the globe moves us toward a more predictable framework that will better allow us to understand future biodiversity loss – and how we might mitigate it – given the expected future expansion of urban areas.

## Acknowledgments

We thank Ellie Diamant for her insights at the conception of the study. JXW was supported by the National Science Foundation Graduate Research Fellowship DGE-2034835.

## Supporting Information

We tested for the effects of the difference in spatial scale between VIIRS and eBird checklists. The VIIRS night-time lights imagery has a spatial resolution of ∼500 m, much smaller than the 5 km filter applied to eBird checklists, and so a single point value may not be representative of the landscape sampled during a specific checklist. In order to check whether this affected our index, we experimented on six example cities, one from each continent (Los Angeles, Buenos Aires, London, Nairobi, Mumbai, Sydney). For every checklist locality within each city, we sampled 100 points from a bivariate normal distribution of latitude and longitude centered on the checklist locality, with a standard deviation of 1 km, truncating values >5 km from the locality. This sampling approach created a scatter of points around the locality, from which a mean radiance value can be calculated. From these mean estimates, we then calculated the mean radiance values for each species across localities. We found that these species-level estimates based on sampled points were highly correlated to the estimates based on single radiance values per locality (r = 0.97–0.98; Fig. S3). Thus, our Urban Association Index (UAI) for each species is the mean radiance value across records where the radiance value of each record is taken from a single pixel of radiance.

**Figure S1.**
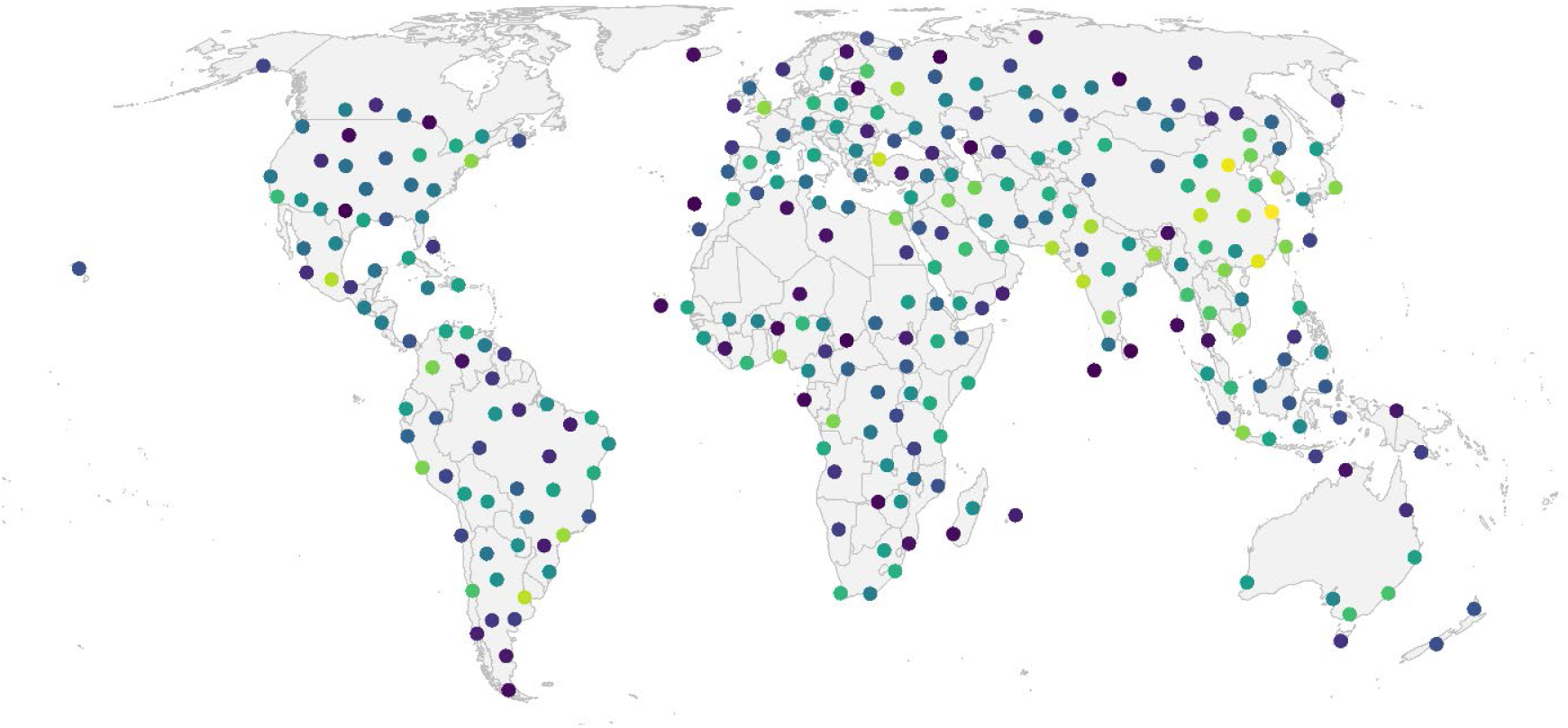
A map of 289 cities spaced ≥500 km apart with populations ≥100,000. Points are colored from least (purple) to most (yellow) populous.

**Figure S2.**
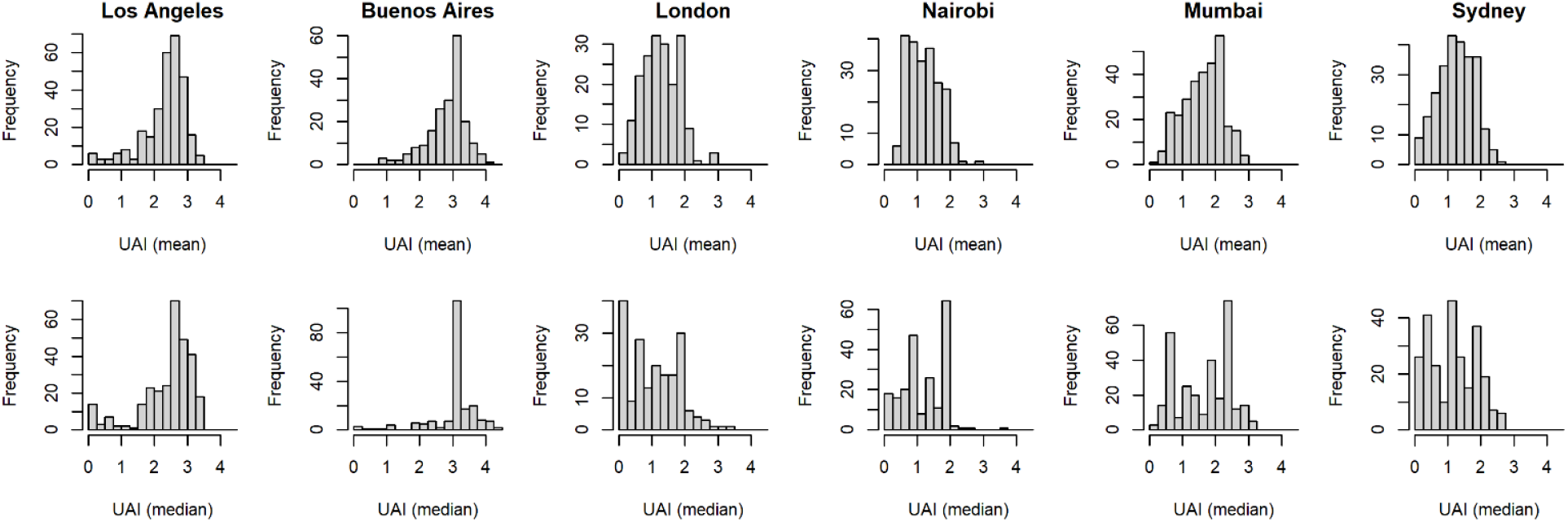
Histograms showing the distribution of Urban Association Indices (UAI) for six example cities. The top row shows UAI estimates based on the mean of radiance values while the bottom row shows UAI estimates based on the median of radiance values.

**Figure S3.**
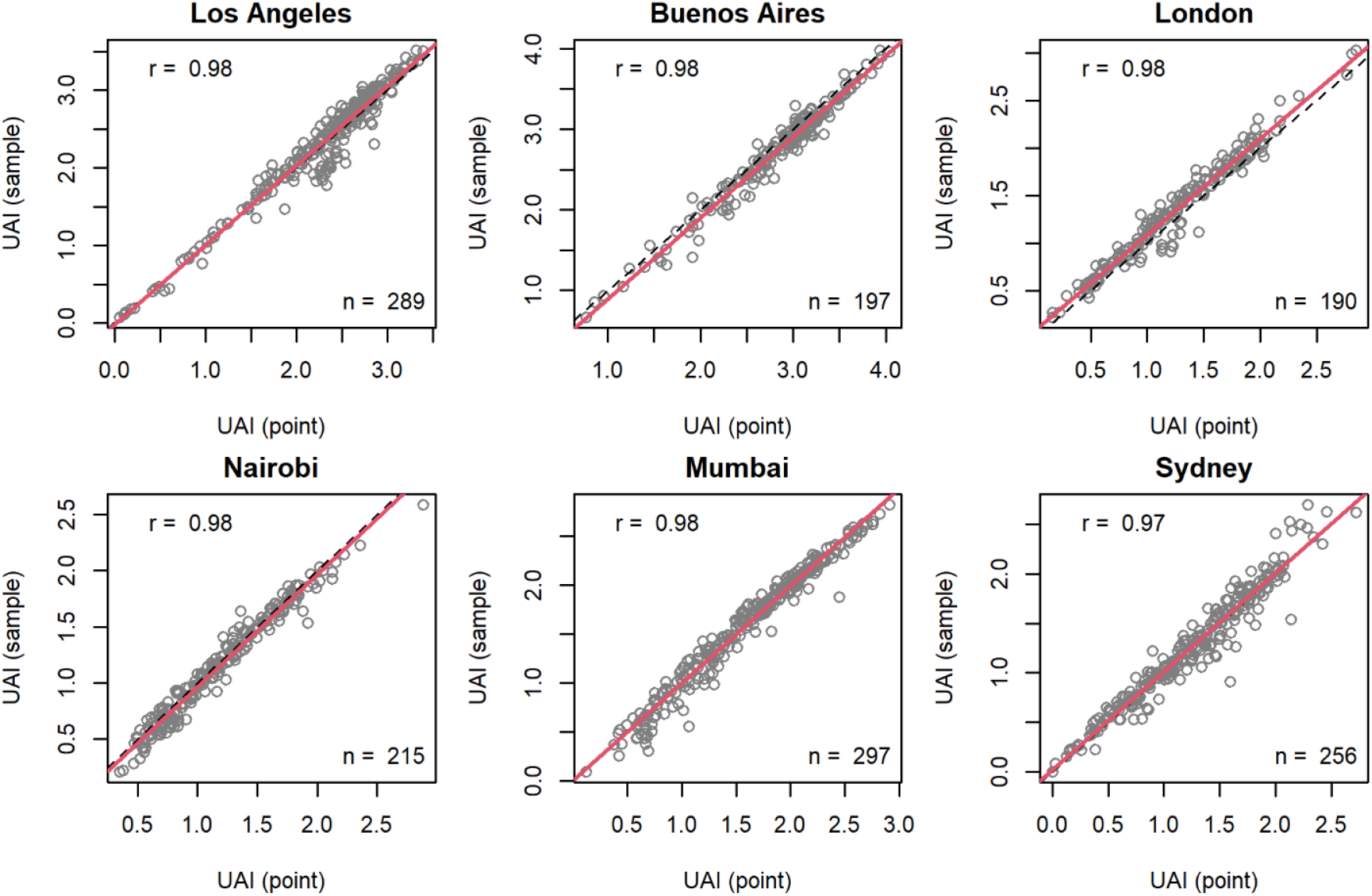
A comparison of two Urban Association Indices (UAI) for six example cities. UAI (point) estimates are based on radiance values from a single pixel of night time lights for each locality. UAI (sample) estimates are based on the mean of 100 points sampled randomly from around each locality. Each point represents the UAI of a species. The dashed line shows 1:1 correspondence, the red line shows the trend between the two indices, and the Pearson’s correlation coefficient is given in the top-right corner.

**Figure S4.**
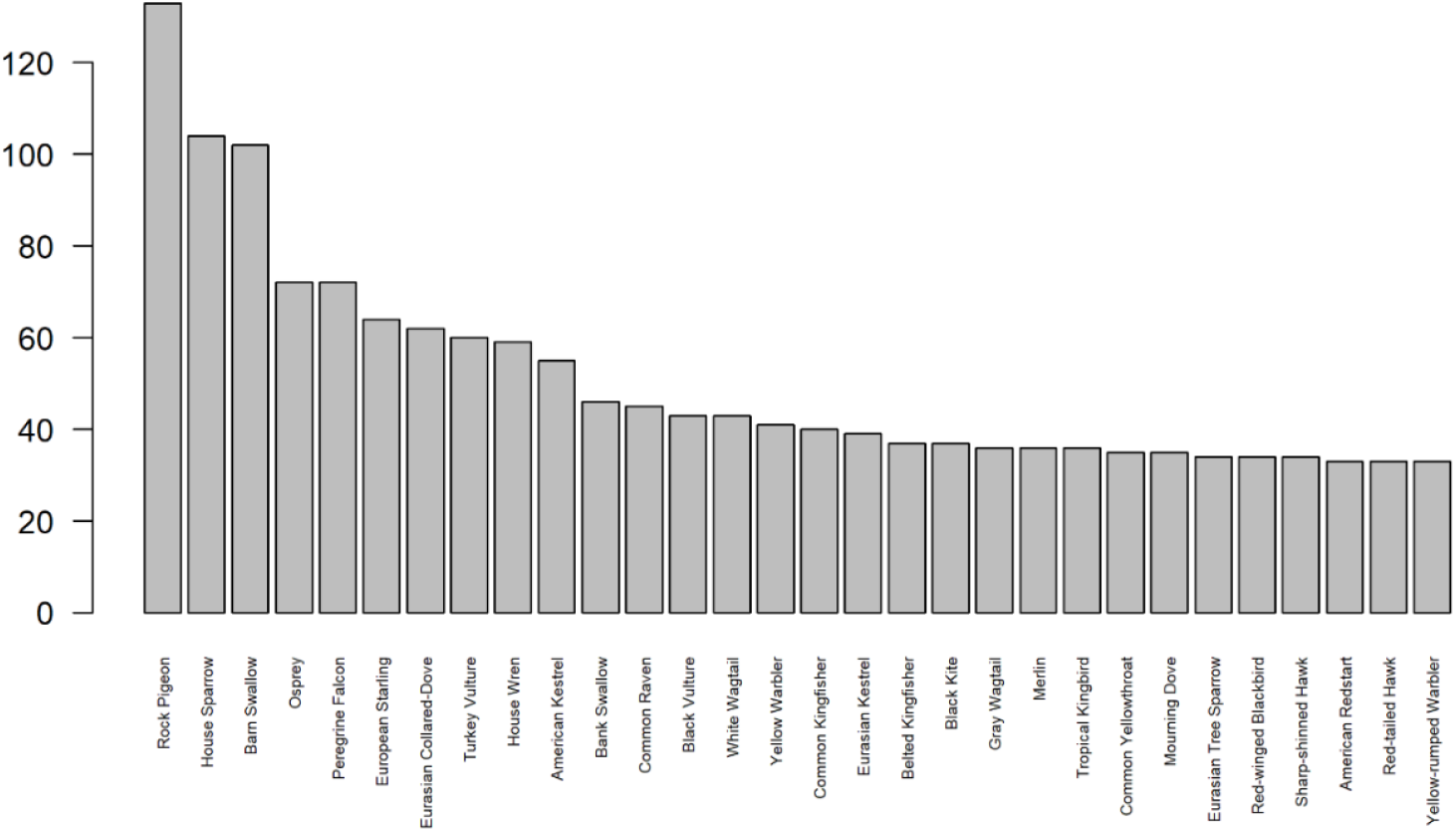
The 50 species represented in the most city circles.

**Figure S5.**
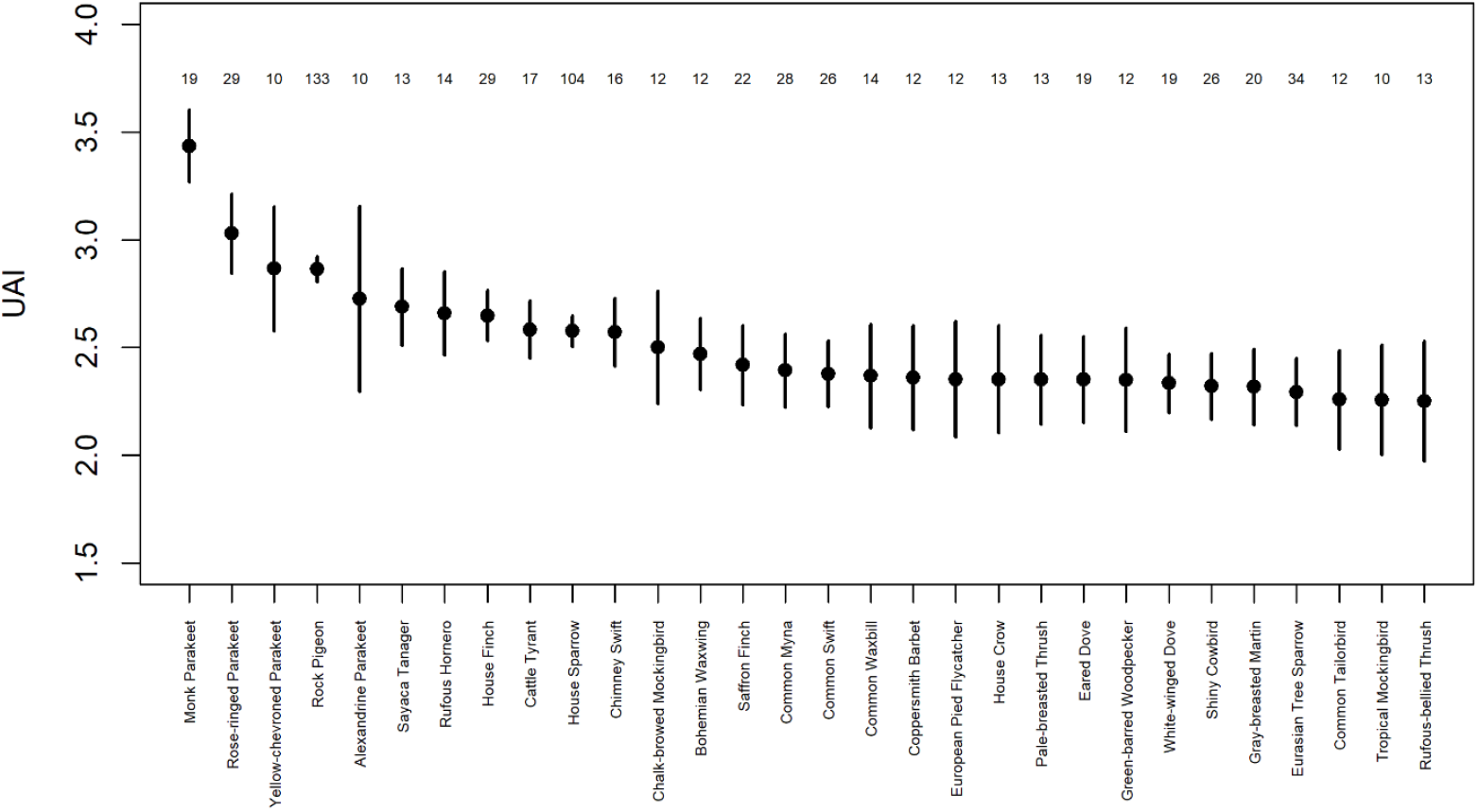
The 30 species with the highest Urban Association Index (UAI). For each species the mean and standard error of their UAI is shown, along with the sample size (i.e., number of cities). Only species present across ≥10 city circles are shown.

**Figure S6.**
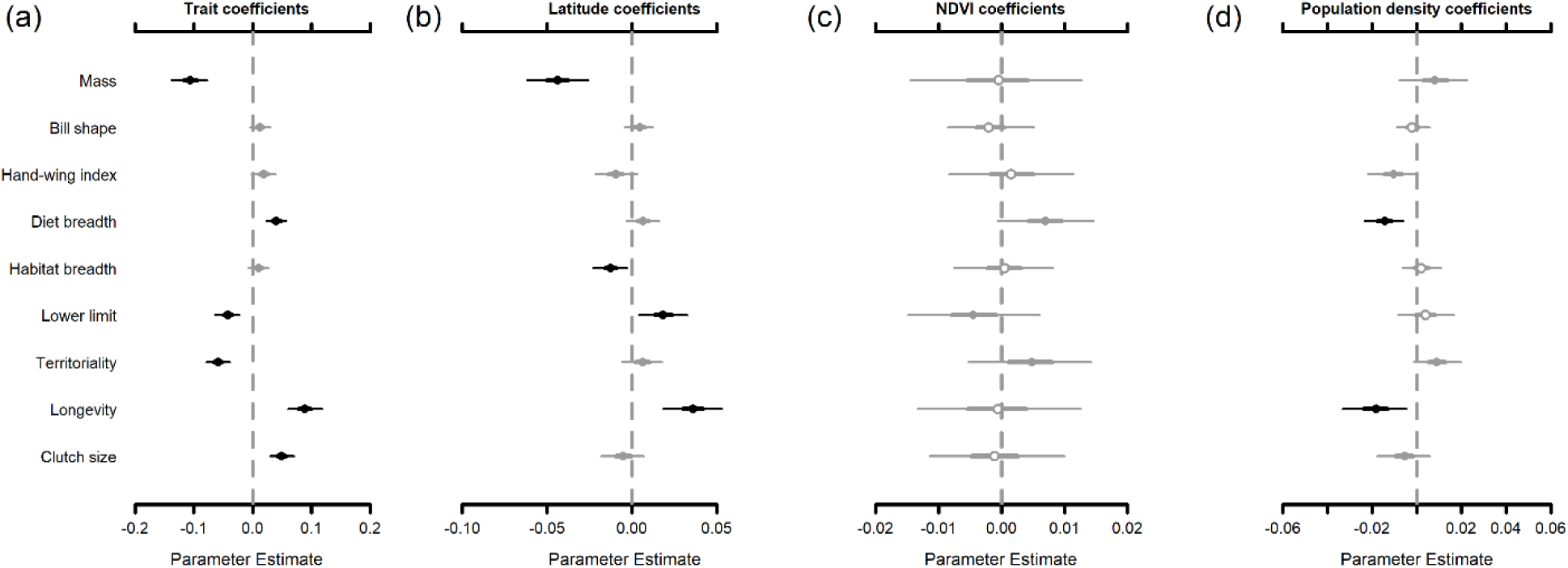
The relationship between Urban Association Index (UAI), species traits, and city covariates for the 920 birds species each represented in at least five cities. This figure is comparable with the results for the full species set in Fig. 4. Covariate coefficients (*β*) show how UAI varies as a function of nine numerical trait covariates (a). In turn, these trait coefficients vary across cities as a function of three city-level variables with corresponding coefficients (*θ*): (b) latitude, (c) NDVI, and (d) human population density. Points show the covariate coefficient estimates with corresponding interquartile range (thick lines) and 95% credible intervals (thin lines). Points are open when the interquartile range overlaps 0. Points and lines are gray when the 95% credible intervals overlap 0 and black when they do not.

